# SFPQ intron retention, reduced expression and aggregate formation in central nervous system tissue are pathological features of amyotrophic lateral sclerosis

**DOI:** 10.1101/2020.09.22.309062

**Authors:** Alison L. Hogan, Natalie Grima, Jennifer A. Fifita, Emily P. McCann, Benjamin Heng, Sandrine Chan Moi Fat, Ram Maharjan, Amy K Cain, Lyndal Henden, Ingrid Tarr, Katharine Y. Zang, Qiongyi Zhao, Zong-Hong Zhang, Amanda Wright, Sharlynn Wu, Marco Morsch, Shu Yang, Kelly L. Williams, Ian P. Blair

## Abstract

**Background:** Splicing factor proline and glutamine rich (SFPQ, also known as polypyrimidine tract-binding protein-associated-splicing factor, PSF) is a RNA-DNA binding protein with roles in key cellular pathways such as DNA transcription and repair, RNA processing and paraspeckle formation. Dysregulation of SFPQ is emerging as a common pathological feature of multiple neurodegenerative diseases including amyotrophic lateral sclerosis (ALS). Increased retention of *SFPQ* intron nine and nuclear loss of the protein have been linked to multiple genetic subtypes of ALS. Consequently, SFPQ dysregulation has been hypothesised to be a common pathological feature of this highly heterogeneous disease.

**Methods:** This study provides a comprehensive assessment of SFPQ pathology in large ALS patient cohorts. *SFPQ* gene expression and intron nine retention were examined in multiple neuroanatomical regions and blood from ALS patients and control individuals using RNA sequencing (RNA-Seq) and quantitative PCR (RT-qPCR). SFPQ protein levels were assessed by immunoblotting of patient and control motor cortex and SFPQ expression pattern was examined by immunofluorescent staining of patient and control spinal cord sections. Finally, whole-genome sequencing data from a large cohort of sporadic ALS patients was analysed for genetic variation in *SFPQ*.

**Results:** *SFPQ* intron nine retention was significantly increased in ALS patient motor cortex. Total *SFPQ* mRNA expression was significantly downregulated in ALS patient motor cortex but not ALS patient blood, indicating tissue specific *SFPQ* dysregulation. At the protein level, nuclear expression of SFPQ in both control and patient spinal motor neurons was highly variable and nuclear depletion of SFPQ was not a consistent feature in our ALS cohort. However, we did observe SFPQ-positive cytoplasmic ubiquitinated protein aggregates in ALS spinal motor neurons. In addition, our genetic screen of ALS patients identified two novel, and two rare sequence variants in *SFPQ* not previously reported in ALS.

**Conclusions:** This study shows that dysregulation of SFPQ is a feature of ALS patient central nervous system tissue. These findings confirm SFPQ pathology as a feature of ALS and indicate that investigations into the functional consequences of this pathology will provide insight into the biology of ALS.

## Background

Amyotrophic lateral sclerosis (ALS) is characterised by the rapid degeneration of motor neurons leading to progressive paralysis and death, typically within 3-5 years of symptom onset (1). ALS is linked clinically, pathologically and genetically with a form of dementia – frontotemporal dementia (FTD), with the two diseases considered to lie on a spectrum of neurodegenerative disease (2). Approximately 10% of ALS patients have a known family history of the disease. Disease-causal mutations have been identified in over 20 genes, which function through a variety of cellular processes (1). In addition to genetic heterogeneity, ALS shows significant clinical variability including age of onset, clinical presentation and rate of progression (3). Disease heterogeneity presents a significant challenge to efforts to unravel disease pathobiology and identify effective therapeutics.

While ALS is a heterogeneous disease, patients share a common pathological feature - the presence of ubiquitinated protein aggregates within affected motor neurons. The majority of patients also show TDP-43 pathology, characterised by cytoplasmic mislocalisation and aggregation of TDP-43 (4,5). TDP-43 pathology is common to sporadic (SALS) and familial (FALS) ALS patients with the exception of genetic subtypes who carry pathogenic mutations in *SOD1* (4) or *FUS* (6,7). Recent evidence suggests that dysregulation of SFPQ may similarly be a pathological feature of multiple subtypes of ALS, including cases without TDP-43 pathology (8).

SFPQ is a predominantly nuclear protein with a range of functions required for cell development and survival, including DNA repair, transcriptional regulation, post-transcriptional RNA processing, paraspeckle formation and axonal transport (9,10). Dysregulation of SFPQ has been linked to multiple neurodegenerative diseases. Altered expression level and loss of nuclear expression, has been reported in animal models and small case-control studies of Alzheimer’s disease (AD) and frontotemporal dementia (FTD) (11–13) and altered methylation of *SFPQ* has been reported in a digenic mouse model of Parkinson’s disease (PD) (14).

In studies of *SFPQ* dysregulation in ALS, increased retention of *SFPQ* intron nine was demonstrated in neural precursor cells derived from fibroblasts of familial patients with ALS-linked mutations in *VCP, SOD1* and *FUS* (8). Whether this pathology is present in mature motor neurons of ALS patients has not been established. Nuclear clearance of SFPQ has been reported in induced pluripotent stem cell (iPSC)-derived motor neurons generated from ALS patient fibroblasts (8), in the motor neurons of a transgenic pig model of ALS (TDP-43^M337V^) (11) and two mouse models of ALS (SOD1^G93A^, VCP^A232E^) (8). However, studies of SFPQ nuclear expression in ALS patient post-mortem tissue have produced inconsistent findings. Significant nuclear clearance of SFPQ was reported in three SALS patients compared to controls (8), while no loss of nuclear SFPQ was reported in two FALS patients with a *FUS* mutation (15). Both studies relied on small patient cohorts and their conflicting findings indicate a need for large cohorts to clarify SFPQ nuclear expression in ALS patient motor neurons and to investigate a potential association between loss of nuclear SFPQ and genetic or pathological subtypes of ALS.

Genetic variation in *SFPQ* has been linked to ALS through the identification of two novel sequence variants in *SFPQ* in FALS patients (16) located in adjacent amino acids within a domain responsible for SFPQ localisation, paraspeckle formation and transcriptional regulation (17). Neither variant was able to rescue motor neuron deficits in a *SFPQ* null mutant zebrafish model, suggesting an impairment of SFPQ function (16). However, segregation with disease could not be tested in either case, thus a definitive causal link between the variants and ALS is yet to be established.

We report analysis of SFPQ pathology in large ALS patient cohorts and multiple disease relevant tissues, including brain and spinal cord. *SFPQ* intron nine retention, SFPQ expression at the mRNA and protein levels and SFPQ protein localisation and aggregation were examined. Our analysis confirmed that intron nine retention was increased in ALS patient motor cortex and demonstrated that SFPQ was a component of the ubiquitinated protein aggregates characteristic of ALS pathology. We also identified two novel and two rare *SFPQ* variants in SALS patients not previously reported. Collectively, our data suggest that SFPQ dysregulation is a significant pathological feature of ALS patient tissue. Aberrant SFPQ may offer a new avenue to explore the mechanisms of ALS and investigate novel therapeutic targets and disease biomarkers that are applicable to the majority of patients.

## Methods

### Study design

This study assessed SFPQ pathology in ALS patient samples at the mRNA and protein levels and screened whole-genome sequencing data from SALS patients to identify genetic variants in *SFPQ. SFPQ* gene expression and the incidence of *SFPQ* intron nine retention was examined in multiple brain regions of ALS patients and control individuals using a combination of RNA-seq and RT-qPCR. *SFPQ* gene expression was also examined in peripheral blood using RNA-seq data from a large ALS case-control cohort. SFPQ protein expression was examined by Western blot analysis of patient cortex and subcellular localisation of SFPQ was examined by immunofluorescent (IF) staining of spinal cord sections from ALS patients with different genetic diagnoses and pathologies. Whole-genome sequencing data from a large cohort of Australian SALS patients was also interrogated to identify novel variants in *SFPQ*.

#### Participants

The cohorts used in this study, totalling 819 participants, are outlined in each section below. This study was approved by the human research ethics committee of Macquarie University (5201600387). Peripheral blood DNA and RNA samples from ALS patients and unrelated controls were obtained from the Macquarie University Neurodegenerative Diseases Biobank and the Australian MND DNA bank. Fresh-frozen and formalin-fixed paraffin-embedded tissues were obtained from the New South Wales Brain Bank Network (Sydney Brain Bank at Neuroscience Research Australia and the New South Wales Brain Tissue Resource Centre at the University of Sydney).

### SFPQ intron retention and transcript expression

#### Cohort details

Two case-control cohorts were used for the analysis of *SFPQ* intron nine retention and *SFPQ* gene expression. The first cohort (CNS-RNA cohort) was used to assess *SFPQ* in RNA extracted from the central nervous system, including the motor cortex, frontal cortex, cerebellum and hippocampus. This cohort comprised 18 ALS cases and 12 controls (details in Supplementary Table 1). The second cohort (Blood-RNA cohort), was used to assess *SFPQ* in RNA extracted from peripheral blood. This cohort comprised 30 SALS cases and 27 controls (details in Supplementary Table 2).

#### RNA extraction

Total RNA was extracted from fresh-frozen tissue from four brain regions (motor cortex, frontal cortex, hippocampus, and cerebellum) using the AllPrep DNA/RNA Mini kit (Qiagen, Germany) according to manufacturer’s instructions, including optional on-column DNase treatment. Total RNA was extracted from peripheral blood using the QIAsymphony PAXgene blood RNA kit and QIAsymphony SP automated instrument (Qiagen, Germany). RNA concentration was measured using the QIAxpert system (Qiagen, Germany) and RNA integrity number (RIN) was determined by Agilent RNA 6000 Nano assay on the Agilent 2100 Bioanalyzer system (Agilent Technologies, USA). Only samples with RIN ≥7 were used for analysis.

#### RNA-seq library preparation, sequencing and pre-processing quality control

A subset of matched samples from the CNS-RNA cohort (six ALS and four control) and the whole Blood-RNA cohort underwent RNA sequencing. RNA-seq libraries were prepared from 1 μg of total RNA using the TruSeq Stranded mRNA LT Sample Prep kit (Illumina, USA). Sequencing was performed on the Illumina NovaSeq 6000 (CNS-RNA-seq cohort subset) or HiSeq2000 (Blood-RNA-seq cohort) platform. The quality of raw sequencing reads was evaluated using fastQC (v0.11.7) for both datasets (18). Trimming and alignment was performed using either Trimmomatic (v. 0.38) (19) or Cutadapt (v1.8.1) (20) respectively and HISAT2 (v2.0.5 and v2.1.0 respectively) (21).

#### RNA-seq data processing and expression analysis

For the CNS-RNA cohort, StringTie (v1.3.4) (22) was used to assemble alignments into gene transcripts. The fragment per kilobase of transcript per million mapped reads (FPKM) of all *SFPQ* mRNA transcripts was calculated for each sample. To examine total *SFPQ* expression, the FPKM of all *SFPQ* transcripts were combined and a two-tailed Mann Whitney t-test was performed to examine difference between cases and controls. To examine intron nine retention, the FPKM of intron nine positive transcripts was normalised to total expression of all transcripts and a two-tailed t-test was performed to compare the difference between cases and controls.

For the Blood-RNA cohort, all data processing and analysis was completed in R (v3.6.2), using BioConductor package edgeR (v. 3.28.1) (23). A standard edgeR trimmed mean of M values normalisation and filtering (filterByExpr) pipeline was used in data processing with 11616 genes remaining for analysis. Counts per million (cpm) for every gene were calculated using normalised library sizes for each sample. Welch two-sample t-test was performed to compare the difference in peripheral blood expression of *SFPQ* in cases and controls.

#### Reverse transcription and RT-qPCR analysis of motor cortex RNA

For quantitative PCR (RT-qPCR), RNA extracted from the motor cortex of the CNS-RNA cohort were analysed. Reverse transcription of 500 ng of motor cortex RNA was performed using the Tetro cDNA Synthesis kit (Bioline, Meridian Bioscience, USA) with random hexamer primers according to the manufacturer’s instructions. RT-qPCR was performed using TaqMan Fast Advanced Master Mix (ThermoFisher Scientific, USA) and the Applied Biosystems ViiA 7 Real-time PCR System (ThermoFisher Scientific, USA). TaqMan assays (ThermoFisher Scientific, USA) were used to measure gene expression of *SFPQ* (Hs00915444_m1) and three reference genes: *B2M* (Hs99999907_m1), *GAPDH* (Hs99999905_m1) and *UBC* (Hs00824723_m1) determined to be appropriate for use as reference genes by qbase+ software program v 3.2 (24). A custom TaqMan assay was designed to assess levels of *SFPQ* intron nine retention (one primer and probe located in intron nine and one primer in exon 10; sequence not disclosed by manufacturer). Cycle threshold (Ct) values were corrected for singleplex or multiplex assay amplification efficiencies. Details of TaqMan assays are provided in Supplementary Table 3.

Gene and transcript expression levels were calculated using the ΔΔCt method, normalised to all three reference genes. *SFPQ* intron nine retention was expressed as the ratio of *SFPQ* intron nine positive mRNA to total *SFPQ* gene expression as previously described (25). Statistical analysis of expression data from cases and controls was performed with two-tailed, unpaired t-tests using the Holm-Sidak method.

### SFPQ protein analysis in patient tissue

#### Cohort details

Fresh-frozen motor cortex samples and formalin-fixed paraffin-embedded cervical spinal cord was obtained for analysis of SFPQ protein expression (CNS protein cohort, details provided in Supplementary Table 4). Motor cortex samples were available for 15 ALS cases and 4 controls, all of which underwent Western blot analysis. Spinal cord sections were available for the whole cohort (20 ALS cases and 7 controls). These samples underwent immunofluorescent (IF) staining.

#### Protein collection from motor cortex tissue

Detergent (RIPA) soluble protein fractions were extracted from motor cortex tissue. Sections were homogenized in 5X volume (μL/mg) of RIPA buffer (50 mM Tris, 150 mM NaCl, 1% Triton-X-100, 5 mM EDTA, 0.5% sodium deoxycholate, 0.1% SDS, pH 8.0) containing phosphatase and protease inhibitors (Roche, Switzerland) using a motor-driven pestle. Homogenates were centrifuged at 124,500 x g for 40 minutes at 4°C and the supernatant was collected (RIPA-soluble fraction). Total protein concentration was determined using the Pierce BCA Protein Assay Kit (ThermoFisher Scientific).

#### Western blot analysis

Protein lysates were prepared in dH2O with NuPAGE LDS sample buffer and reducing agent (ThermoFisher Scientific, USA) and denatured at 70°C for 10 minutes. Protein lysates were electrophoresed on NuPAGE 4 – 12% Bis-Tris gels (ThermoFisher Scientific, USA) in NuPAGE MOPS SDS buffer supplemented with NuPAGE antioxidant (ThermoFisher Scientific, USA). Protein was transferred to Immobilon-FL PVDF membrane (Merck, USA) using a wet transfer system (Bio-Rad, Criterion Blotter). Membranes were blocked in Odyssey Blocking Buffer in TBS (LI-COR Biosciences, USA) for 1 hour at room temperature followed by overnight incubation at 4°C with primary antibodies: 0.6 μg/mL polyclonal rabbit anti-SFPQ (ab38148, Abcam, UK), and 1:5000 monoclonal mouse anti-GAPDH (60004-1-Ig, Proteintech, USA). Membranes were then incubated for 1 hour at room temperature with IRDye 800CW donkey anti-rabbit IgG and 680LT donkey anti-mouse IgG, 1:20,000 (LI-COR Biosciences). Membranes were visualised using the Odyssey CLx imaging system and protein bands quantified with the Image Studio Lite software (LI-COR Biosciences).

#### Immunofluorescent staining of spinal cord tissue

Spinal cord tissue sections were heated at 70 °C for 30 minutes, deparaffinised with xylene and rehydrated with a descending series of ethanol and water. Antigen retrieval was performed by heating the sections in 10 mM citrate buffer (pH 6.0, Sigma-Aldrich, USA) at > 96 °C for 20 minutes. Sections were washed in PBS and blocked with 5% normal goat serum (NGS, Vector Laboratories, USA) with 0.1% TWEEN 20 in PBS for 1 hour at room temperature. Primary antibody incubation was performed overnight at 4°C. Primary antibodies used were 1:100 rabbit anti-SFPQ (Abcam), 1:5000 monoclonal mouse anti-TDP-43 phosphorylated Ser409/410 (TIP-PTD-M01, Cosmo Bio, Japan), 1:150 monoclonal mouse anti-ubiquitin (MAB1510, Merk Millipore, MA, US). Following PBS washes, sections were incubated with goat anti-rabbit and goat anti-mouse secondary antibodies conjugated to Alexa Fluor 555 or 488 (1:250, Life Technologies) for 1 hour at room temperature. Sections were then incubated in NeuroTrace 640/660 Deep-Red Fluorescent Nissl Stain (1:100, Life Technologies) for 20 minutes at room temperature and mounted with ProLong Gold Antifade Mountant with DAPI (Life Technologies).

Sections were imaged with a ZEISS LSM 880 inverted confocal laser-scanning microscope. Quantification of the intensity of SPFQ expression in the nucleus and the cytoplasm of Nissl stained ventral horn motor neurons was performed using the free drawing tool and the Measure function in FIJI-Image J software. Fluorescence intensity of SFPQ in the nucleus was divided by intensity in the cytoplasm to give the nuclear cytoplasmic ratio (N:C). All neurons identified within a section (minimum of ten) were analysed. Statistical analysis was performed with one-way ANOVA with Kruskall-Wallis test for multiple comparisons.

### Genetic analysis of SFPQ

#### SFPQ variant identification

Truseq PCR-free whole-genome sequencing (WGS) data from 609 SALS cases were used for genetic analysis (26). Custom UNIX scripts were applied to identify all variants in the *SFPQ* gene (NM_005066). For comparison, the non-neurological subset of non-Finnish Europeans (nNFE, n=51,592) from the Genome Aggregation Database (GnomAD) (27) was used as a control cohort. Filtering was applied to identify all novel and rare protein-altering (missense, frameshift and non-frameshift insertions/deletions, stop gain/loss, splicing) and 3’ and 5’ untranslated region (UTR) variants. Variants with a minor allele frequency of > 0.0001 in the GnomAD nNFE control cohort were excluded.

*In silico* protein prediction analyses were used to evaluate the potential pathogenic role of *SFPQ* variants in ALS. Prediction annotations were applied to each variant using dbNSFP v3.3a (28) and were scored as benign or pathogenic (29). Twenty-one programs were used to predict functional consequences and conservation across species of each variant (PolyPhen2-HDIV and -HVAR, LRT, MutationTaster, Mutation Assessor, FATHMM, PROVEAN, VEST3, MetaSVM, MetaLR, M-CAP, CADD, DANN, FATHMM-MKL, Eigen, GenoCanyon, fitscons, GERP++, phyloP, phastCons, and SiPhy). Each variant was also screened through four additional ALS patient datasets (Project MinE (n= 4366, (30)), ALS data browser (ALSdb; http://alsdb.org/, n=3093), ALS variant server (AVS; http://als.umassmed.edu/, n=1415) and study accession phs000101.v5.p1 (n=247) from the database of Genotypes and Phenotypes (dbGAP)).

#### Gene burden analysis

Fisher’s exact tests were applied in R to determine if *SFPQ* carried a burden of rare qualifying protein-altering or UTR variants in SALS cases compared to the gnomAD nNFE control cohort. The minor allele frequencies of <0.005 and <0.0001 were used to identify qualifying rare variants in the SALS and GnomAD nNFE cohorts respectively. As 3’UTR and 5’UTR variants do not alter protein sequence but may affect gene expression, the two types of variation were analyzed separately. A Bonferroni corrected significance threshold of p<0.025 (n=2) was applied.

#### Variant association analysis

Fisher’s exact tests were applied in R to identify potential ALS risk or protective variants, by comparing major and minor allele counts of variants between SALS cases and the gnomAD nNFE control cohort. All biallelic *SFPQ* variants excluding intergenic variants were analysed, and a Bonferroni corrected significance threshold of 3.33×10^−4^ was used to account for the 150 variants under analysis.

### Statistical analyses

All statistical analyses were performed using either GraphPad Prism (Prism v8 software, GraphPad) or R v3.6.2 (R Foundation for Statistical Computing, Vienna, Austria, 2018 https://www.r-project.org/). *P* values equal to or less than 0.05 were considered to be statistically significant, except for genetic analyses where Bonferroni corrections for multiple testing were applied.

## Results

### Cohort analysis

Analysis of the case-control cohorts used to assess *SFPQ* expression, intron retention, SFPQ localisation and aggregate formation is summarised in **Table 1**. No significant differences in the age of onset, sex or post-mortem interval were present between cases and controls with the exception of the motor cortex RT-qPCR cohort, in which differences in age of death approached significance (*p* = 0.05). We therefore assessed the potential effect of age of death on RNA quality (and RT-qPCR analysis) using linear regression analysis. No correlation between age of death and RNA quality (determined by RIN) was observed (*R* = 0.041, *p* = 0.83, supplementary Figure 1).

**Table 1:** Statistical comparisons of case-control cohorts used to assess *SFPQ* expression, intron retention, SFPQ localisation and aggregate formation.

| Analysis | Cohort | Sample type | n (% female) | Case-control comparison |
| --- | --- | --- | --- | --- |
| RNA-seq | CNS-RNA (subset of) | Motor cortex (MC), Frontal cortex (FC), Hippocampus (HP), Cerebellum (CB) | 6 SALS<sup>a</sup> (50%)<br>4 controls<sup>b</sup> (50%) | Age: $p=0.832$<br>Sex: $p=1$<br>PMI: $p=0.338$<br>RIN MC: $p=0.207$<br>RIN FC: $p=0.406$<br>RIN HP: $p=0.978$<br>RIN CB: $p=0.888$ |
| RNA-seq | Blood-RNA | Peripheral blood | 30 SALS<sup>c</sup> (53%)<br>27 controls (47%) | Age: $p=0.318$<br>Sex: $p=0.317$<br>RIN: $p=0.442$ |
| RT-qPCR | CNS-RNA (subset of) | Motor cortex | 17 ALS (59%)<br>12 controls (54%) | Age: $p=0.05^*$<br>Sex: $p=0.682$<br>PMI: $p=0.608$<br>RIN: $p=0.751$ |
| Western blot | CNS protein | Motor cortex | 15 ALS (50%)<br>4 controls<sup>d</sup> (30%) | Age: $p=0.548$<br>Sex: $p=0.589^e$<br>PMI: $p=0.849$ |
| IF staining | CNS protein | Cervical spinal cord | 20 ALS<sup>f</sup> (50%)<br>7 controls<sup>d,f</sup> (30%) | Age: $p=0.548$<br>Sex: $p=0.589^d$<br>PMI: $p=0.849$ |

PMI – post-mortem interval, RIN – RNA integrity number
\* statistically significant difference between cases and controls
<sup>a</sup> five of six ALS cases were also analysed through RT-qPCR
<sup>b</sup> all controls were also analysed through RT-qPCR
<sup>c</sup> six SALS are also present in the CNS RNA cohort
<sup>d</sup> two control samples are also present in the CNS RNA cohort
<sup>e</sup> Chi-squared approximation may be incorrect due to $n<5$ in one category
<sup>f</sup> 15 ALS and 4 controls were also analysed through Western Blot

### SFPQ intron nine retention in motor cortex, hippocampus, frontal cortex and cerebellum

To determine whether increased *SFPQ* intron nine retention is a feature of ALS patient CNS tissue, we analysed RNA-seq data from RNA extracted from four brain regions (n = 6 ALS, n = 4 controls): motor cortex (the most severely affected brain region in ALS), frontal cortex and hippocampus (regions variably affected in ALS) and cerebellum (region largely spared from ALS pathology). Transcript analysis of RNA-seq data identified 12 *SFPQ* transcripts, four of which retained intron nine (**Fig. 1a**). All 12 transcripts were present in ALS patients and controls in all brain regions. The percentage of intron nine retaining transcripts was consistently elevated in the three affected brain regions of ALS patients compared to controls; motor cortex (1.2 fold greater), frontal cortex (2.5 fold greater) and hippocampus (1.7 fold greater) but did not reach significance (**Fig. 1b**). No increase in intron nine retention was evident in the cerebellum of ALS patents compared to controls.

**Figure 1.**
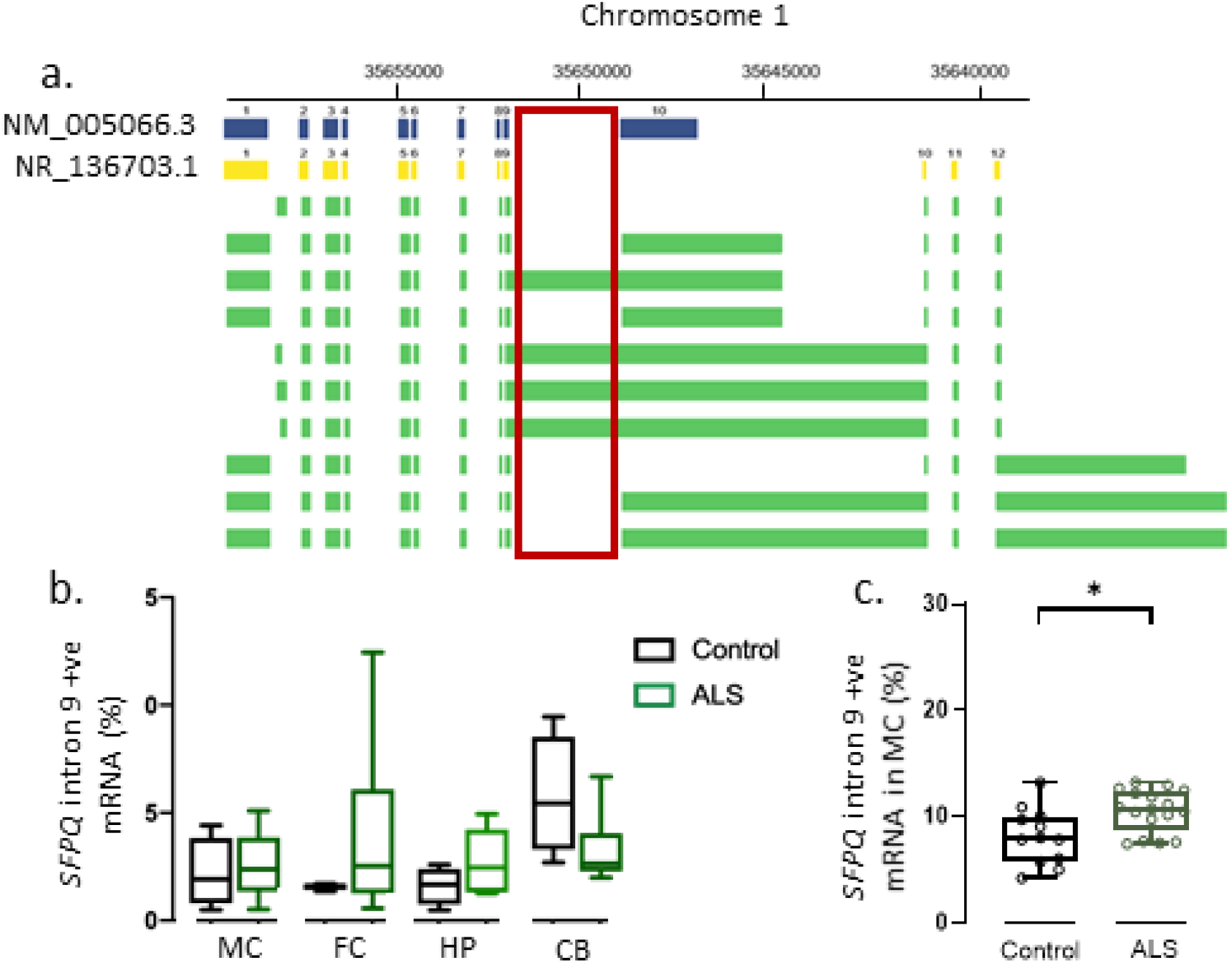
RNA-seq and RT-qPCR analysis of *SFPQ* intron retention in motor cortex of the ALS patient-control cohort. **a**. Twelve *SFPQ* transcripts were present in RNA-seq data from four brain regions of six ALS patients and four controls. Two transcripts aligned with established SFPQ RNA sequences (NM_005066.3, NR_136703.1). Four of 12 transcripts retained intron nine (red box). **b**. Quantification of *SFPQ* intron nine positive (+ve) transcripts relative to total *SFPQ* in the motor cortex (MC), frontal cortex (FC), hippocampus (HP) and cerebellum (CB). No significant increase in intron nine positive transcripts was observed in any of the four brain regions examined between ALS patients and controls. **c**. RT-qPCR analysis of motor cortex (MC) RNA in the cortical cohort (n = 17 cases, n = 12 controls). A significant increase in the relative number of intron nine positive transcripts was identified in ALS patients compared to controls (*p* = 0.019).

RT-qPCR was used to further investigate *SFPQ* intron nine retention in the motor cortex of an extended cohort (n = 17 SALS, n = 12 controls). This revealed the intron nine positive transcripts relative to total *SFPQ* transcripts to be significantly higher in ALS patients compared to controls (*p* = 0.006, **Fig 1c)**.

### SFPQ gene expression in central nervous system and blood

To investigate *SFPQ* gene expression in the CNS, we examined RNA-seq data for each of the four brain regions and the RT-qPCR analysis of the motor cortex. From the RNA-seq data, *SFPQ* gene expression was found to be lower in ALS patient motor cortex (*p* = 0.01), frontal cortex (*p* = 0.01), hippocampus (*p* = 0.02) and cerebellum (*p* = 0.01) compared to controls (**Fig. 2a**). RT-qPCR analysis also demonstrated significantly lower *SFPQ* expression in ALS patient motor cortex compared to controls (*p* = 0.019) (**Fig. 2b**).

**Figure 2.**
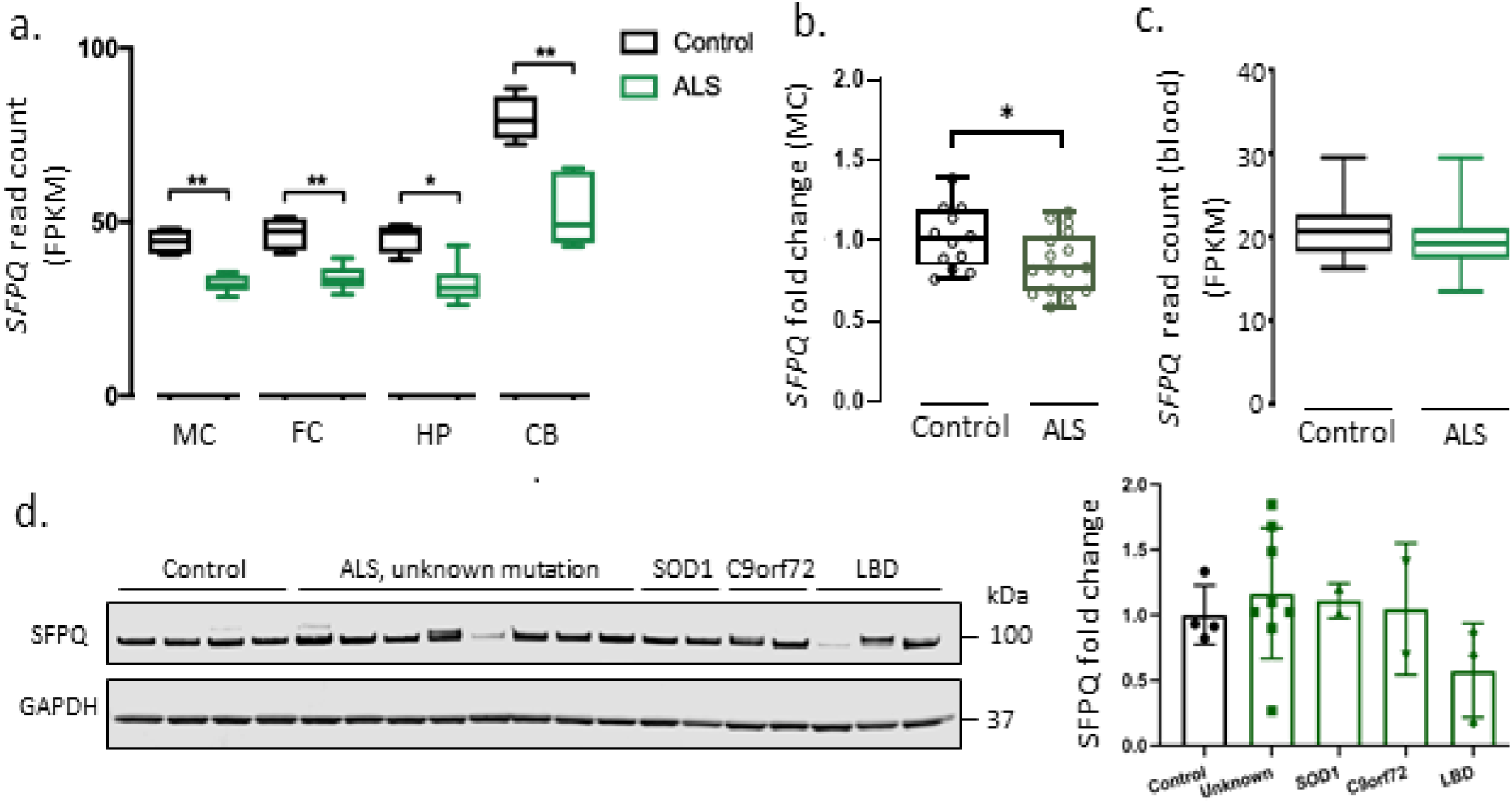
Total *SFPQ* gene expression is reduced in ALS patient motor cortex, frontal cortex, hippocampus and cerebellum, but not in blood or at the protein level. **a**. Expression of *SFPQ* was quantified in motor cortex (MC), frontal cortex (FC), hippocampus (HP), and cerebellum (CB) by RNA-seq (n = 4 controls, n= 6 ALS patients). A significant reduction in total *SPFQ* expression was observed between controls and ALS cases in the MC, FC, CB and HP. **b**. Total *SFPQ* expression was examined in the motor cortex (MC) of the extended cortical cohort (n = 17 cases, n = 12 controls) through RT-qPCR. Significantly reduced *SFPQ* expression was observed in ALS patients compared to controls (*p* = 0.0255**) c**. *SFPQ* expression in blood was assessed by RNA-seq in a separate cohort (n = 30 patients, n = 22 controls). No significant difference in *SFPQ* expression was observed. **d**. Western blot analysis of SFPQ protein expression in the motor cortex of controls (n = 4) and ALS patients (n = 8 unknown mutation, n = 2 SOD1, n = 2 C9orf72, n=3 with lewy body pathology, LBD). SFPQ expression was normalised to GAPDH loading control. SFPQ expression was variable between individuals, however no significant difference between patients and controls was found.

To investigate whether the reduced *SFPQ* gene expression observed in ALS patient brain was generalised or tissue specific, *SFPQ* expression was examined in a RNA-seq dataset of peripheral blood collected from 30 patients and 27 controls with RIN values ≥7 (Blood-RNAseq cohort). No difference in *SFPQ* FPKM was observed in peripheral blood between patients and controls (*p* = 0.2) (**Fig 2c**).

### SFPQ protein expression in motor cortex

To investigate whether the observed reduction in *SFPQ* mRNA expression was translated to SFPQ protein expression, Western blot analysis of patient motor cortex tissue was performed on detergent (RIPA) soluble lysates collected from a cohort of 15 ALS patients and four age-matched controls. SFPQ protein expression varied between individuals, however no consistent difference in SFPQ expression was observed in ALS patients compared to controls (**Fig. 2d**). A subset of ALS patients who demonstrated Lewy body pathology in addition to TDP-43 pathology (n = 3) demonstrated the lowest SFPQ expression of all groups examined (1.74 fold lower than control individuals).

### SFPQ localisation in human spinal motor neurons

We next investigated SFPQ expression in spinal motor neurons of 20 ALS patients from different genetic and pathological subtypes of ALS, including two *SOD1* cases, seven *C9orf72* cases and 11 cases with unknown mutation including three cases with Lewy body pathology in addition to TDP-43 pathology. Strong nuclear expression of SFPQ was observed in the majority of neurons in both patients and controls. However, neurons with equivalent SFPQ expression in the nucleus and cytoplasm, as well as neurons with complete nuclear depletion of SFPQ were observed. Representative images of the SFPQ localisation phenotypes are shown in **Fig 3a**.

**Figure 3.**
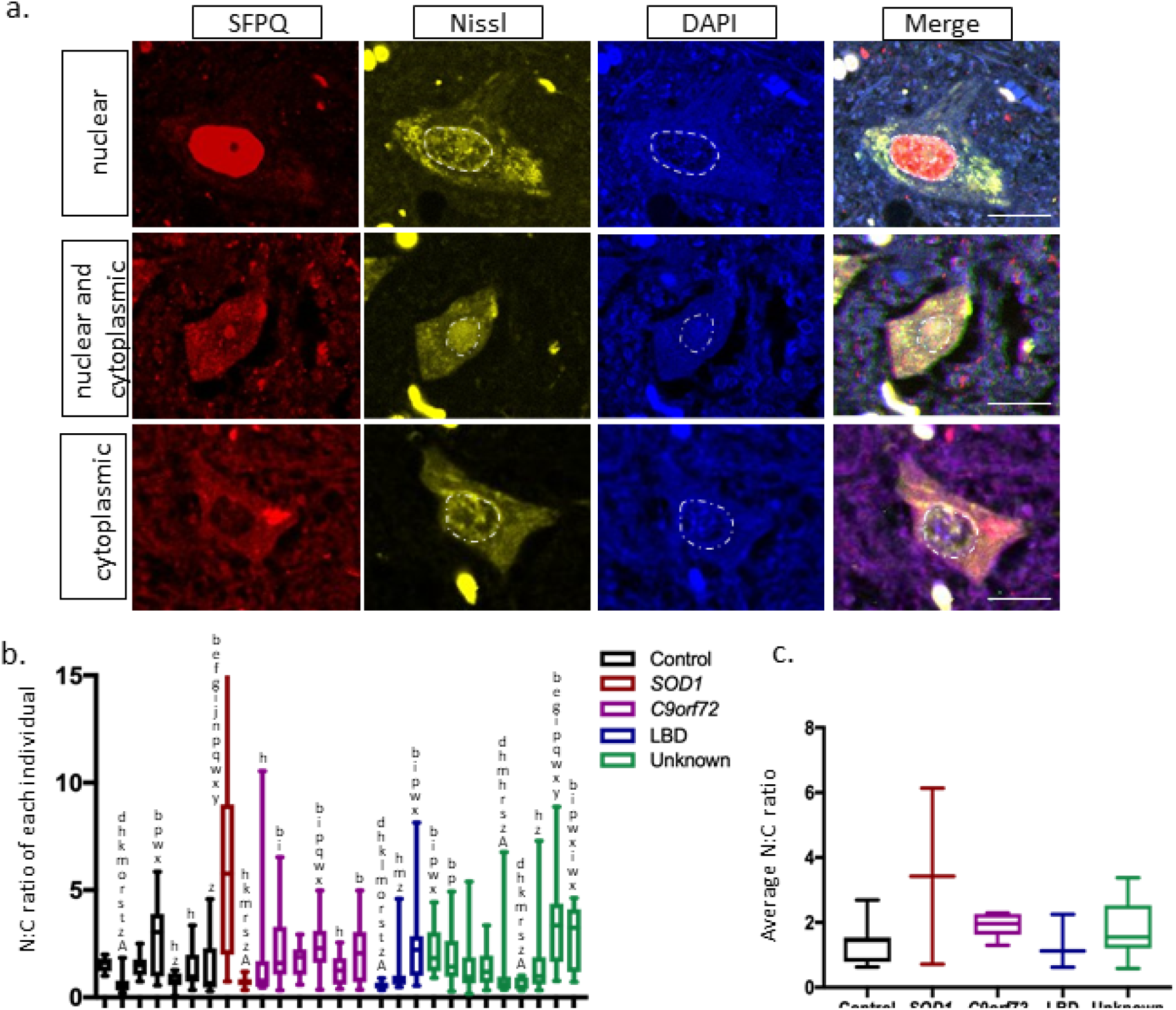
Subcellular localization of SFPQ in spinal motor neurons. **a**. Representative images of the three SFPQ localisation phenotypes observed in human spinal motor neurons: strong nuclear expression, equal nuclear and cytoplasmic expression and nuclear depletion with low cytoplasmic expression. Strong nuclear expression was the most common phenotype observed. However, all three phenotypes were observed in ALS patients and controls. Scale bar: 20 µm. **b**. Quantification of SFPQ nuclear cytoplasmic ratio (N:C) in all motor neurons identified for each individual (n = 10 - 30) demonstrated significant differences in SPFQ subcellular localisation between individuals. However, significant differences between individuals did not associate with disease or mutation status. Letters indicate individuals in which a significant difference in N:C ratio was observed. **c**. The average N:C ratio of each individual did not differ between controls and any ALS genetic or pathological subgroup examined.

To quantify nuclear expression of SFPQ relative to cytoplasmic expression, the SFPQ nuclear cytoplasmic ratio was determined for every neuron identified in all samples (n = 10 – 29 neurons per individual). Statistically significant variability in SFPQ nuclear cytoplasmic ratio was observed between individuals in both the control and patient groups (**Fig 3b**). Within the control group, average nuclear cytoplasmic ratio ranged from 0.46 to 2.69, with significant differences between individuals up to *p* = 0.0002. Similar variability was evident within ALS patient subgroups, including between the two patients with a *SOD1* mutation (*p* <0.0001), the *C9orf72* patients (up to *p* = 0.009), patients with unknown genetic causes (up to *p* = 0.0001) and patients with Lewy body pathology (up to *p* = 0.001). As a result of this intra-group variability, no significant difference in SFPQ N:C ratios were observed between controls and ALS patients, or between different genetic or pathological subtypes of ALS (**Fig 3c**).

### SFPQ positive aggregates in ALS patient neurons

Immunofluorescent (IF) staining of the cohort with SFPQ and pTDP-43 antibodies identified protein aggregates in 68 spinal motor neurons from 14 ALS patients (**Fig 4A**). Of the 68 neurons, 18 contained aggregates positive for both pTDP-43 and SFPQ (26.5 %), three neurons contained SFPQ-positive pTDP-43-negative aggregates (4.4 %) and 47 neurons contained SFPQ-negative pTDP-43-positive aggregates (69.1 %) (**Fig 4C**). SFPQ positive aggregates were identified in two *C9orf72* patients, three patients with an unknown mutation and one patient with Lewy body pathology. No SFPQ aggregates were observed in controls or patients with a *SOD1* mutation. Patients with SFPQ or TDP-43 aggregates in their motor neurons did not show a reduction in average SFPQ nuclear cytoplasmic ratio compared to controls or compared to ALS patients without neuronal aggregates (**Fig.4d)**.

**Figure 4.**
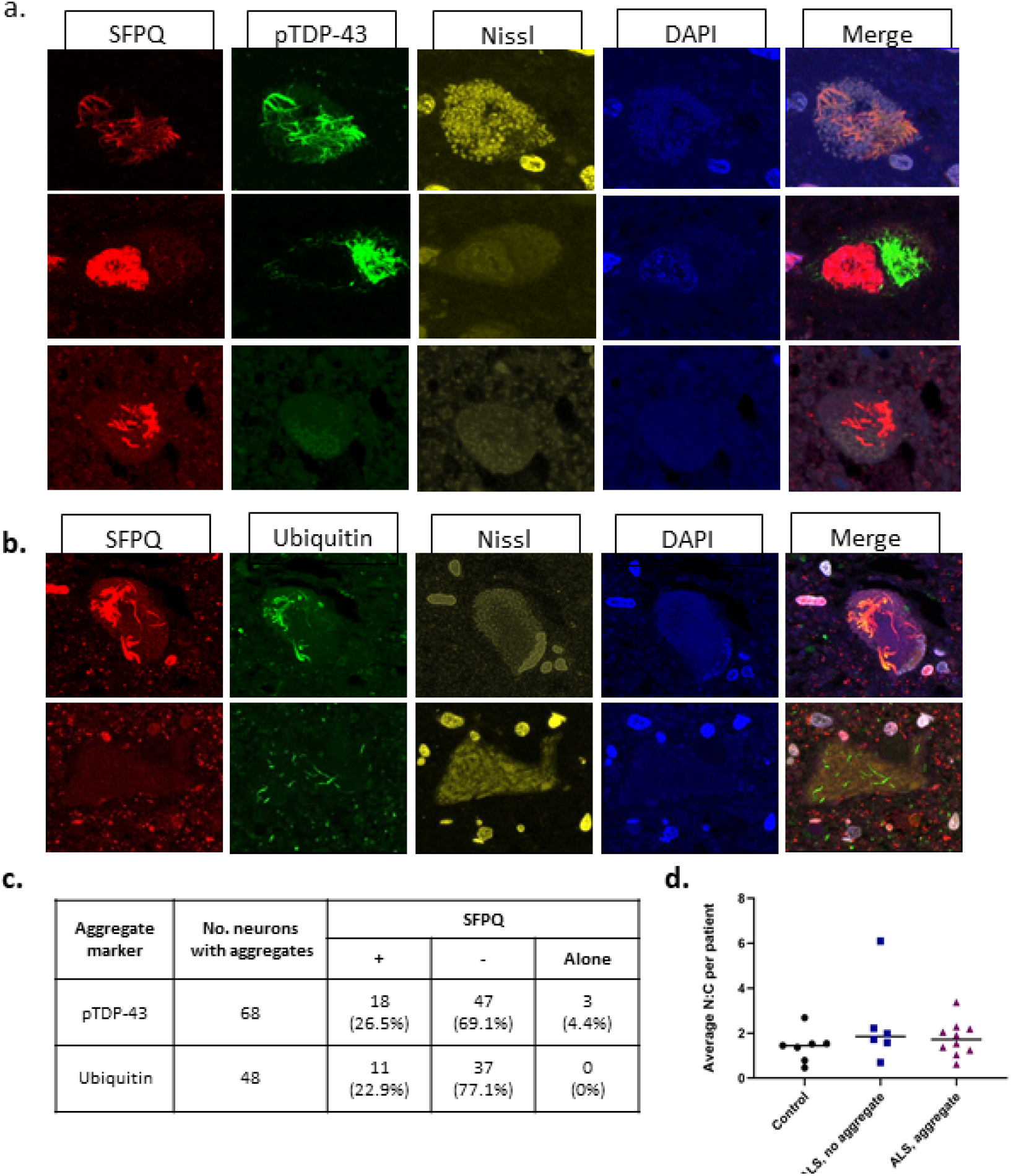
SFPQ-positive aggregates were identified in spinal motor neurons of ALS patients. **a**. Spinal cord sections immunostained with SFPQ and pTDP-43 identified neurons with pTDP-43- and SFPQ-positive aggregates (top row) pTDP-43-positive SFPQ-negative aggregates with stron SFPQ nuclear expression (middle) and SFPQ-positive, pTDP-43-negative aggregates (bottom row). **b**. Spinal cord sections immunostained with SFPQ and ubiquitin identified neurons with ubiquitin-positive and SFPQ-positive aggregates and ubiquitin-positive SFPQ-negative aggregates. No SFPQ-positive ubiquitin-negative aggregates were identified. **c**. pTDP-43-positive aggregates were more commonly found in ALS patient neurons than SFPQ-positive aggregates. Co-aggregation of SFPQ and TDP-43 was observed in ∼ 27.5% of cases. All SFPQ aggregates were ubiquitinated, but not all ubiquitin-positive aggregates were SFPQ positive. **d**. No relationship between the presence of aggregates and SFPQ cytoplasmic accumulation (changes in N:C ratio) was evident.

To determine whether SFPQ positive aggregates were ubiquitinated, additional sections from a subset of the patient cohort (10 patients known to carry SFPQ or TDP-43 positive aggregates) were IF stained with SFPQ and ubiquitin antibodies. Ubiquitinated aggregates were identified in 48 neurons from the 10 samples, 11 of which were positive for SPFQ (22.9%) (**Fig 4b, 4c)**. No ubiquitin negative SFPQ positive aggregates were identified.

### SFPQ sequence variants in sporadic ALS patients

Two novel *SFPQ* variants were previously reported in FALS patients (NM_005066, c.1597A>C, p.N533H and c.1600C>A, p.L534I) including an Australian case from our familial cohort (31). In this study, we interrogated a large cohort of 609 Australian SALS cases to determine whether the same, or novel *SFPQ* variants were present. Two additional novel variants, a non-synonymous variant (c.G436C, p.G146R), and a non-frameshift deletion (c.812_814delTCT, p.K271_272del) absent from GnomAD nNFE control dataset, and two rare non-synonymous point variants c.C2015T, p.A672V (rs371481157) and c.C229T, p.P77S (rs763571943) (present in one and two individuals in the GnomAD nNFE dataset, minor allele frequency of < 0.0001) were identified, one in each of four different SALS individuals (Supplementary Table 5). The rare p.P77S variant was also observed in an ALS patient from the Project MinE dataset. The novel p.G146R and rare p.P77S variants were predicted to be benign by the majority of prediction tools, and the rare p.A672V variant was predicted pathogenic by 13 of 17 prediction tools and is located at an amino acid residue highly conserved across species (Supplementary Table 5). MutationTaster (32) and PROVEAN (33) predicted the novel deletion to be disease-causing and neutral, respectively.

An enrichment of rare *SFPQ* protein-altering variants was seen among SALS patients (1.31%) compared to controls (0.40%; Fisher’s exact p= 0.00444), as well as an enrichment of *SFPQ* UTR variants in SALS patients (1.64%) compared to controls (0.41%; p= 0.0004797). A total of 150 biallelic variants were identified in *SFPQ* and underwent variant association testing. None of the variants were associated with SALS in this cohort.

## Discussion

### Overview

This study identified increased *SFPQ* intron nine retention, reduced *SFPQ* gene expression and SFPQ protein aggregation as pathological features of ALS patient CNS tissue. SFPQ is a multifunctional protein with regulatory roles in numerous cellular pathways that are disrupted in ALS, including transcriptional regulation, post-transcriptional processing, DNA repair and paraspeckle formation (10,34). Due to these essential functions, dysregulation of SFPQ is predicted to have wide-ranging downstream effects. Indeed, loss of SFPQ function is associated with embryonic death in mice and zebrafish (16,35,36). In this study, we characterised SFPQ pathology in ALS patients and identified novel and rare *SFPQ* gene variants in SALS cases. These data further implicate SFPQ dysregulation in ALS.

### Increased retention of *SFPQ* intron nine in CNS tissue is a feature of ALS

Increased retention of *SFPQ* intron nine has previously been reported in immature motor neurons derived from ALS patient fibroblasts (8). Here, we analysed ALS post-mortem brain tissue and confirmed that increased *SFPQ* intron nine retention is a pathological feature of ALS motor cortex. Interestingly, while it did not reach statistical significance, SFPQ intron nine retention was also elevated in the frontal cortex and hippocampus, regions that are secondarily affected in ALS. Frontal cortex and hippocampal pathology are primary features of FTD and AD (37), neurodegenerative diseases in which dysregulation of SFPQ at the protein level has been reported. Findings form this study suggest that an analysis of *SFPQ* intron nine retention may be warranted in these disorders.

Intron retention can have multiple consequences including alternate protein isoforms with novel or altered function, altered subcellular localisation and transcriptional regulation, and induction of nonsense mediated decay (reviewed in 9,10). *In situ* hybridisation studies in iPSCs derived from ALS patients has demonstrated increased intron nine-positive *SFPQ* mRNA in the cytoplasm relative to the nucleus (8). Further, analysis of iCLIP and eCLIP data from cell lines has shown that the SFPQ protein (8) and FUS (40) bind extensively to *SFPQ* intron 9, findings that have lead the hypothesis that loss of nuclear SFPQ and FUS (a feature of a subset of ALS patients), is a consequence of *SFPQ* intron nine retention pathology (40). Further examination of the *SFPQ* intron nine interactome in ALS-relevant cells may offer novel insights into protein mislocalisation in ALS.

*SFPQ* intron nine carries 46 stop codons, the first of which occurs at codon seven, suggesting if intron nine positive transcripts are translated (rather than undergoing nonsense mediated decay), a truncated isoform will be produced. The putative truncated isoform would lack the C-terminal canonical nuclear localisation sequence (one of two nuclear localisation sequences), potentially altering SFPQ subcellular localisation. It is yet to be determined whether this isoform can be detected in ALS patient tissue. The biological consequences of nuclear loss of SFPQ, including potential loss- or gain-of-function, also awaits further study.

### SFPQ gene expression is reduced in CNS tissue of ALS cases

Both our RNA-seq and RT-qPCR analyses demonstrated that *SFPQ* gene expression was reduced in multiple brain regions of ALS patients. However, this reduced expression was not evident in peripheral blood, suggesting a brain-specific *SFPQ* pathology. The significance of this pathology however is unclear. SFPQ protein levels, as determined by mass spectroscopy, have previously been shown to be unaltered in ALS patient frontal cortex (41), a finding supported by our Western blot analysis of ALS patient motor cortex. However, further investigation is warranted. Our analysis used non-overlapping sample cohorts for *SFPQ* transcript and protein analyses, preventing us from performing a direct correlation between *SFPQ* mRNA and protein expression.

While a decrease in SFPQ protein was not observed across the whole ALS cohort, expression was 1.74 fold lower in the subset of ALS patients with Lewy body pathology compared to control individuals. ALS with Lewy body pathology is rare (42,43) and may form an as-of-yet poorly characterised pathological subset of ALS. SFPQ pathologies specific to clinical subsets of AD (44) and FTD (45) patients have been described, including downregulation of SFPQ expression and nuclear depletion. Further examination of SFPQ in additional ALS patient cohorts may identify distinct clinical subgroups with reduced expression.

### SFPQ localisation was not consistently altered in spinal cord neurons

Nuclear loss of SFPQ has been demonstrated in iPSC-derived motor neurons (8) and multiple animal models that overexpress ALS-linked transgenes (8,11). However, this change in SFPQ expression pattern has not been consistently demonstrated in patient tissue, either in previous reports (8,15) or the current study. Two previous studies in small cohorts (n=3) presented conflicting reports of loss of nuclear SFPQ expression (8,15), while the current study, in a larger case control cohort, demonstrated that SFPQ expression pattern varies significantly between individuals. Interestingly, marked fluctuation in SFPQ nuclear expression is associated with the circadian rhythm (46), suggesting that the variability between individuals seen in this study may, in part, reflect time of death.

By using post-mortem tissue, we have demonstrated that increased *SFPQ* intron nine retention and decreased *SFPQ* gene expression are features of end-stage disease. It remains possible that nuclear loss of SFPQ occurs at an earlier point in disease and that neurons that develop this pathology are lost prior to post-mortem. Animal models provide the opportunity to investigate this hypothesis as they allow analysis of early-to-mid stage disease pathology and examination of pathology at consistent timepoints, thereby eliminating circadian rhythm- or patient-specific variability.

### SFPQ-positive aggregates were observed in ALS patient motor neurons

This is the first report of SFPQ-positive aggregates in the motor neurons of ALS patients. These aggregates were found in spinal cord motor neurons of patients with a pathogenic mutation in *C9orf72* and in patients who did not carry a mutation in a known ALS-linked gene. No SFPQ-positive aggregates were observed in the *SOD1* cases. Patients who carry *SOD1* mutations account for less than 2% of ALS cases (47) and display a distinct pathology characterised by SOD1 positive aggregates that are negative for TDP-43. While the number of *SOD1* patients in this study was small, our findings suggest that SFPQ-positive cytoplasmic aggregates are not a feature of SOD1 pathology, but rather are a feature of the TDP-43 pathology that is characteristic of the majority of FALS and SALS patients.

SFPQ is an obligate dimer - to perform its many functions it must efficiently bind to other proteins (17,34). SFPQ contains a highly charged coiled-coil domain essential for polymerisation (17), and like all RNA binding proteins, an intrinsically disordered, low complexity region that affects the folding state and solubility of the protein (often referred to as prion-like domain). While the ability to functionally aggregate is essential for SFPQ function, it does give rise to a propensity for pathological aggregation under stress conditions (48). It will be important to determine whether SFPQ is passively sequestered in pre-formed aggregates or whether it can act as a scaffold for protein aggregation, actively contributing to the formation of protein aggregates.

### *SFPQ* variants in ALS patients

The identification of SFPQ positive aggregates in patient neurons supports the hypothesis of a genetic link between *SFPQ* and ALS. Of the proteins identified in ubiquitinated aggregates within ALS patient neurons, most have been found to carry ALS-linked mutations. This includes *TARDBP* (4), *FUS* (6,7), *OPTN* (49), *UBQLN2* (50,51) and *C9ORF72* (52,53). In healthy control individuals, *SFPQ* demonstrates little genetic variance, which indicates a low tolerance for variation (1) and in turn, suggests protein-altering *SFPQ* variants are likely to have pathogenic implications. Two missense variants that alter adjacent residues in the SFPQ coiled-coil domain (c.1597A>C, p.N533H and c.1600C>A, p.L534I) have previously been reported in FALS patients (16). We screened *SFPQ* for both these, and other novel gene variants, in a large cohort of SALS patients. The previously reported variants were not present, nor were any additional variants within the coiled-coil domain. However, two additional novel and two rare variants were identified, one of which (p.A672V) is predicted to have a pathogenic effect by the majority of *in silico* analysis tools. In addition, an enrichment of rare protein-altering and UTR variants was present in SALS cases compared to controls. Taken together, these results suggest a role for genetic variation of *SFPQ* in ALS, including causative gene mutations (16) and variants of low disease effect that may act as risk factors to developing disease or influence ALS phenotypes. Further analysis of additional patient cohorts and functional analyses of identified novel and rare *SFPQ* variants is required to determine their pathogenic role as causative mutations or risk factors in ALS.

### Conclusions

We present the first report of SFPQ pathology in large ALS cohorts and confirm that SFPQ dysregulation in the form of increased retention of intron nine, reduced gene expression and the formation of ubiquitinated aggregates are pathological features of ALS. Investigation of the pathological and mechanistic consequence of this dysregulation will provide insight into the biology of ALS and other neurodegenerative diseases.

