## Supplementary figures and images for "SFPQ intron retention, reduced expression and aggregate formation in central nervous system tissue are pathological features of amyotrophic lateral sclerosis"

### Supplementary figure 1

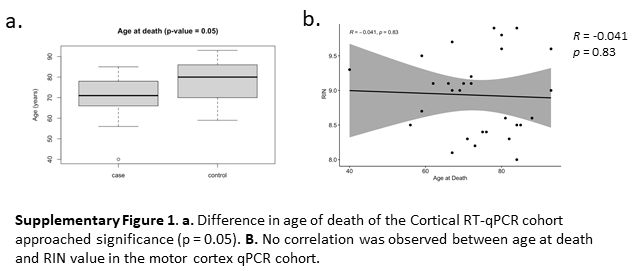
