## Supplementary table 1 for "SFPQ intron retention, reduced expression and aggregate formation in central nervous system tissue are pathological features of amyotrophic lateral sclerosis"

Supplementary table 1. ALS case and control central nervous system RNA (CNS-RNA) cohort used for RNA-seq and RT-qPCR analysis.

| **ID #** | **Status** | **Age** | **Sex** | **PMI** | **pH** | **MC RIN** | **CB RIN** | **FC RIN** | **HP RIN** |
| --- | --- | --- | --- | --- | --- | --- | --- | --- | --- |
| C1a* | Control | 67 | Male | 29 | 7.06 | 9 | 9.6 | 9.7 | 9.2 |
| C2a* | Control | 67 | Female | 15.5 | 6.42 | 8.1 | 8.9 | 8.5 | 8.1 |
| C3a* | Control | 75 | Male | 34 | 6.77 | 8.4 | 8.6 | 8.4 | 8.7 |
| C4a* | Control | 80 | Female | 29 | 6.32 | 9.8 | 10 | 9.7 | 10 |
| C5a | Control | 73 | Female | 45 | 6.86 | 8.2 | N/A | N/A | N/A |
| C6a | Control | 80 | Male | 12 | 6.5 | 9.6 | N/A | N/A | N/A |
| C7a | Control | 59 | Male | 49 | 6.86 | 9.5 | N/A | N/A | N/A |
| C8a | Control | 93 | Female | 7 | 6.44 | 9.6 | N/A | N/A | N/A |
| C9a | Control | 88 | Female | 31 | 6.18 | 8.6 | N/A | N/A | N/A |
| C10a | Control | 84 | Male | 22 | 5.75 | 8 | N/A | N/A | N/A |
| C11a | Control | 93 | Female | 15 | 6.39 | 9 | N/A | N/A | N/A |
| C12a | Control | 84 | Male | 36 | 6.41 | 9.9 | N/A | N/A | N/A |
| A1a* | ALS | 70 | Female | 5 | 6.41 | 9.1 | 9.3 | 10 | 9.4 |
| A2a* | ALS | 78 | Female | 7 | 6.39 | 9.9 | 10 | 9.5 | 9.6 |
| A3a* | ALS | 72 | Male | 31 | 5.6 | 9.1 | 8.8 | 8.6 | 7.4 |
| A4a* | ALS | 67 | Male | 35 | 6.04 | 9.7 | 8.5 | 9.3 | 9.8 |
| A5a* | ALS | 62 | Female | 8 | 6.25 | 9.1 | 9.7 | 9.8 | 7.9 |
| A6a^ | ALS | 79 | Male | 32 | 6.23 | 9.7 | 9.7 | 9.5 | 9.8 |
| A7a | ALS | 69 | Female | 41 | 6.68 | 9 | N/A | N/A | N/A |
| A8a | ALS | 72 | Female | 8 | 6.43 | 9.2 | N/A | N/A | N/A |
| A9a | ALS | 66 | Male | 24 | 6.4 | 9.1 | N/A | N/A | N/A |
| A10a | ALS | 81 | Female | 19 | 5.94 | 8.6 | N/A | N/A | N/A |
| A11a | ALS | 82 | Female | 24 | 6.27 | 8.3 | N/A | N/A | N/A |
| A12a | ALS | 76 | Male | 15 | 6.21 | 8.4 | N/A | N/A | N/A |
| A13a | ALS | 85 | Female | 24 | 5.8 | 8.5 | N/A | N/A | N/A |
| A14a | ALS | 71 | Female | 25 | 5.97 | 8.3 | N/A | N/A | N/A |
| A15a | ALS | 40 | Female | 57 | 6.42 | 9.3 | N/A | N/A | N/A |
| A16a | ALS | 84 | Male | 8 | 6.24 | 8.5 | N/A | N/A | N/A |
| A17a | ALS | 56 | Male | 60 | 6.12 | 8.5 | N/A | N/A | N/A |
| A18a | ALS | 59 | Female | 20 | 6.31 | 8.7 | N/A | N/A | N/A |

PMI = post-mortem interval (hours), RIN = RNA integrity number, MC = motor cortex, CB = cerebellum, FC = frontal cortex, HP = hippocampus, SC = cervical spinal cord.

29/30 MC samples were used for RT-qPCR analysis

CB, FC and HP samples underwent RNA-seq only
*MC for these individuals was used for RNA-seq and RT-qPCR analysis

^MC for this individual was used for RNA-seq only
