## Supplementary table 2 for "SFPQ intron retention, reduced expression and aggregate formation in central nervous system tissue are pathological features of amyotrophic lateral sclerosis"

Supplementary table 2. ALS patient and control peripheral blood cohort used for RNA-seq analysis.

| **ID #** | **Status** | **Age at initial collection** | **Sex** | **RIN** |
| --- | --- | --- | --- | --- |
| C1b | Control | 72 | Female | 8.2 |
| C2b | Control | 63 | Female | 7.7 |
| C3b | Control | 66 | Female | 7.8 |
| C4b | Control | 43 | Female | 7.0 |
| C5b | Control | 53 | Male | 7.0 |
| C6b | Control | 69 | Male | 7.8 |
| C7b | Control | 38 | Female | 7.8 |
| C8b | Control | 62 | Female | 7.3 |
| C9b | Control | 56 | Male | 7.9 |
| C10b | Control | 57 | Male | 7.3 |
| C11b | Control | 42 | Female | 7.4 |
| C12b | Control | 47 | Female | 7.2 |
| C13b | Control | 72 | Male | 7.1 |
| C14b | Control | 63 | Female | 7.0 |
| C15b | Control | 60 | Female | 7.4 |
| C16b | Control | 58 | Male | 7.1 |
| C17b | Control | 66 | Female | 7.3 |
| C18b | Control | 42 | Male | 7.5 |
| C19b | Control | 74 | Male | 7.5 |
| C20b | Control | 60 | Female | 7.7 |
| C21b | Control | 66 | Female | 7.2 |
| C22b | Control | 53 | Female | 7.8 |
| C23b | Control | 64 | Male | 7.6 |
| C24b | Control | 52 | Female | 7.3 |
| C25b | Control | 60 | Male | 7.3 |
| C25b | Control | 44 | Female | 7.7 |
| C27b | Control | 60 | Female | 7.7 |
| A1b | ALS | 64 | Male | 7.8 |
| A2b | ALS | 57 | Female | 7.6 |
| A3b | ALS | 79 | Male | 7.8 |
| A4b | ALS | 58 | Female | 7.0 |
| A5b | ALS | 60 | Male | 7.5 |
| A6b | ALS | 71 | Male | 7.1 |
| A7b | ALS | 44 | Male | 7.2 |
| A8b | ALS | 66 | Male | 7.1 |
| A9b | ALS | 76 | Male | 9.1 |
| A10b | ALS | 70 | Male | 8.3 |
| A11b | ALS | 69 | Male | 8.5 |
| A12b | ALS | 65 | Female | 7.0 |
| A13b | ALS | 33 | Female | 7.7 |
| A14b | ALS | 39 | Female | 7.3 |
| A15b | ALS | 65 | Male | 7.8 |
| A16b | ALS | 64 | Female | 7.0 |
| A17b | ALS | 85 | Male | 7.3 |
| A18b | ALS | 45 | Male | 8.1 |
| A19b | ALS | 67 | Male | 7.4 |
| A20b | ALS | 33 | Female | 7.9 |
| A21b | ALS | 36 | Male | 7.7 |
| A22b | ALS | 47 | Male | 7.9 |
| A23b | ALS | 63 | Female | 7.2 |
| A24b | ALS | 57 | Female | 7.0 |
| A25b | ALS | 66 | Male | 7.3 |
| A26b | ALS | 61 | Female | 7.2 |
| A27b | ALS | 55 | Male | 7.4 |
| A28b | ALS | 91 | Male | 7.4 |
| A29b | ALS | 72 | Male | 7.5 |
| A30b | ALS | 75 | Female | 7.3 |
