## Supplementary table 3 for "SFPQ intron retention, reduced expression and aggregate formation in central nervous system tissue are pathological features of amyotrophic lateral sclerosis"

Supplementary table 3. Details of TaqMan assays used for RT-qPCR analysis.

| **Target** | **Assay ID** | **Dye** | **RefSeq** | **Assay location** | **Amplicon length (bp)** | **Singleplex or Multiplex** | **Efficiency** | **GeNorm M value^** |
| --- | --- | --- | --- | --- | --- | --- | --- | --- |
| SFPQ intron 9 | Custom assay | FAM | N/A | ND* | 50-100 ** | Multiplex | 99.423% | N/A |
| SFPQ | Hs00915444_m1 | FAM | NM_005066.2 | 1906 | 66 | Multiplex | 100.683% | N/A |
| B2M | Hs99999907_m1 | VIC | NM_004048.2 | 409 | 75 | Multiplex | 90.243% | 0.56 |
| GAPDH | Hs99999905_m1 | VIC | NM_002046.5 | 243 | 122 | Singleplex | 93.229% | 0.54 |
| UBC | Hs00824723_m1 | VIC | NM_021009.6 | 454 | 71 | Multiplex | 98.51% | 0.49 |

* Not disclosed by manufacturer. Submitted 288 nucleotide sequence for assay design. Genomic position hg38 chr1:35,184,455 - 35,184,742.

** Amplicon confirmed via gel electrophoresis.

^ As calculated by qbase+ (reference).
