## Supplementary table 4 for "SFPQ intron retention, reduced expression and aggregate formation in central nervous system tissue are pathological features of amyotrophic lateral sclerosis"

Supplementary table 4. ALS patient and control cohort used for immunofluorescent staining and Western blot analysis.

| **ID** | **Status** | **Subtype** | **Mutation** | **Age** | **Gender** | **PMI** | **pH** | **WB analysis** |
| --- | --- | --- | --- | --- | --- | --- | --- | --- |
| C1c | Control | N/A | N/A | 62 | Female | 35 | 6.06 | No |
| C2c | Control | N/A | N/A | 59 | Female | 24 | N/A | No |
| C3a* | Control | N/A | N/A | 75 | Male | 34 | 6.77 | No |
| C4c | Control | N/A | N/A | 69 | Male | 13.5 | 6.74 | Yes |
| C5c | Control | N/A | N/A | 37 | Male | 24 | 6.7 | Yes |
| C6a** | Control | N/A | N/A | 80 | Male | 12 | 6.5 | Yes |
| C7c | Control | N/A | N/A | 61 | Male | 30 | 6.69 | Yes |
| A1c | ALS | SALS | *C9orf72* | 65 | Female | 18 | 6.42 | No |
| A2c | ALS | SALS | *C9orf72* | 61 | Male | 26.5 | 6.44 | No |
| A3c | ALS | SALS | *C9orf72* | 64 | Female | 7 | 6.78 | No |
| A4c | ALS | ALS | *C9orf72* | 68 | Male | 16 | 6.67 | No |
| A5c | ALS | SALS | *C9orf72* | 61 | Female | 22 | 6.13 | No |
| A6c | ALS | FALS | *C9orf72* | 60 | Male | 99 | 6.6 | Yes |
| A7c | ALS | FALS | *C9orf72* | 75 | Male | 21.5 | 6.6 | Yes |
| A8c | ALS | PMA (FALS) | *SOD1* | 53 | Female | 23 | 6.78 | Yes |
| A9c | ALS | FALS | *SOD1* | 82 | Male | 22 | 6.28 | Yes |
| A10c | ALS | PMA | Unknown | 74 | Female | 20 | 6.7 | Yes |
| A11c | ALS | ALS | Unknown | 54 | Male | 9 | 6.8 | Yes |
| A12c | ALS | ALS | Unknown | 57 | Female | 5 | 6.72 | Yes |
| A13c | ALS | Lewy body pathology | Unknown | 81 | Male | 70 | 6.59 | Yes |
| A14c | ALS | ALS | Unknown | 84 | Male | 23 | 6.69 | Yes |
| A15c | ALS | FALS | Unknown | 54 | Female | 16 | 6.39 | Yes |
| A16c | ALS | ALS | Unknown | 71 | Male | 16 | 7.04 | Yes |
| A17c | ALS | Lewy body pathology | Unknown | 71 | Male | 12.5 | 6.19 | Yes |
| A18c | ALS | Lewy body pathology | Unknown | 64 | Female | 14 | 6.92 | Yes |
| A19c | ALS | ALS | Unknown | 61 | Female | 20 | 6.64 | Yes |
| A20c | ALS | ALS | Unknown | 77 | Female | 9 | 6.34 | Yes |

ALS = amyotrophic lateral sclerosis, SALS = sporadic ALS, FALS = familial ALS, PMA = progressive muscular atrophy, PMI = post-mortem index (hours), WB = Western blot

*control sample is also in CNS RNA-Seq and motor cortex RT-qPCR cohort
**control sample is also in motor cortex RT-qPCR cohort
