## Supplementary table 5 for "SFPQ intron retention, reduced expression and aggregate formation in central nervous system tissue are pathological features of amyotrophic lateral sclerosis"

Supplementary table 5. Novel and rare SFPQ variants identified in Australian sporadic ALS cases.

| **Position (GRCh37)** | **cDNA change** | **protein change** | **rsID** | **gnomAD nNFE MAF** | **Pathogenic functional predictions (total)** |
| --- | --- | --- | --- | --- | --- |
| Chr1:35650166 | c.C2015T | p.A672V | rs371481157 | 0.00001138 | 13 (17) |
| Chr1:35657836 | c.812_814delTCT | p.K271_272del | rs1266283483 | . | 1 (2) |
| Chr1:35658215 | c.G436C | p.G146R | . | . | 5 (17) |
| Chr1:35658422 | c.C229T | p.P77S | rs763571943 | 0.00005382 | 4 (17) |

rsID = Database of variation (dbSNP) variant identification number, gnomAD nNFE = non-neuronal, non-Finnish European subset of the genome aggregation database, MAF = minor allele frequency
